# Tuning the contractility and deformation modes of active actin-based assemblies in vitro: from 2D active networks to liquid crystal drops

**DOI:** 10.1101/2024.12.04.626882

**Authors:** Samantha Stam, Steven Huntley, Carolyn A. Feigeles, Veronica J. Armstrong, Margaret A. Cheves, Sam Rubin, Kimberly L. Weirich

## Abstract

Actin cytoskeleton-based materials are widely investigated as model cellular materials to elucidate physical mechanisms of cell mechanics, such as shape regulation and force production, as well as intriguing soft polymeric materials. In this method, we detail creating actin-based assemblies *in vitro* using purified protein for fluorescence microscopy studies. We polymerize long actin filaments in a sample chamber and use a polymer depletant to crowd filaments into a 2D-entangled network against a surface passivated with a surfactant layer. Adding skeletal muscle myosin II filaments in the presence of ATP induces contraction of the actin network. By bundling actin filaments with cross-linker, we tune the contractility of the assembly, transitioning from a material that buckles to a material that slides at the microscale. By reducing the length of the actin filaments through co-polymerizing actin in the presence of capping protein, we tune the material from being a 2D network to a liquid crystal. Cross-linking of dispersed short actin filaments results in 3D liquid crystal droplet formation.

## INTRODUCTION

Active, biological materials underly mechanical processes in a variety of physiological processes including intracellular transport, cell migration, cell shape regulation, and biological force generation (1–3). *In vitro* assemblies of cytoskeletal systems, constructed from purified and engineered protein components self-assembled in buffer, are an established tool for characterizing fundamental biophysical and biochemical processes (4–7). These simplified, model systems allow proteins to be studied without the complexity of cellular environments, allowing the roles of individual components to be identified. In addition to supporting fundamental biological studies, *in vitro* cytoskeletal assemblies also are widely developed as experimental systems to investigate liquid crystalline and active or out-of-equilibrium materials (8–15).

Actin is a key biopolymer in cytoskeletal assemblies regulating cell mechanics and dynamics through motor and polymerization-driven forces (3, 16). Fluorescence microscopy experiments allow visualization of actin network structural changes during processes such as motor-driven contraction and actin bundling by cross-linking proteins. Imaging three-dimensional actin networks as they are remodeled by embedded motor proteins has illuminated the role of actin cross-linkers for network contraction and pattern formation (12, 17–19). However, quasi-two-dimensional actin networks overcome challenges in imaging with sufficient spatiotemporal resolution to capture network structural changes. Additionally, high-density, quasi-two-dimensional actin structures are a closer mimic of cellular assemblies such as the dense actin cortex of a cell (20). Strategies to produce quasi-two-dimensional actin structures have included attachment of actin nucleating proteins to a coverslip (21, 22), coupling of actin to a surface through linker molecules (23, 10, 24), formation of dense actin bundles attached to beads at a coverslip surface (25, 26), and concentration of actin filaments at a coverslip surface with molecular crowding agents (10, 27, 28).

Here, we detail a method for creating reproducible model actin-based assemblies, and how to modify it to prepare variations from active networks to quasi 2D liquid crystals and nematic droplets through crowding of actin filaments with various accessory proteins to a passivated coverslip surface. We have previously used similar methods to study formation of phase-separated liquid crystalline droplets of short actin filaments with cross-linker addition (11), changes in myosin II-generated actin network deformation modes (e.g. filament buckling or relative sliding) with filament cross-linking and connectivity (28), differences in myosin II-generated forces on actin bundles with varying spacing and compliance (29), and myosin II transport (30). The use of cross-linking proteins and actin capping protein allows for systematic variation of actin filament length and assembly architecture.

## PROTOCOL

### Proteins and labeling

For the purposes of this protocol, we will assume the researcher begins with stocks of purified proteins, which are fluorescently labeled when appropriate for fluorescence microscopy investigation. These proteins can be purchased or purified and fluorescently labeled in the lab (31–37). In this protocol, we use stocks of proteins that have been frozen in liquid nitrogen and stored at −80°C.

The proteins typically can be labeled through similar methods. The protein can be labeled as a step of purification or from a frozen stock. Thaw frozen protein stocks on ice before proceeding with the labeling. The following is a method for fluorescently labeling skeletal muscle myosin II (myosin) with a maleimide-functionalized fluorophore (adapted from (38)).

### Prepare fluorescently-labeled myosin

1. Start with ∼ 2 mL of myosin at a concentration of 5-10 mg/mL. Note: In this protocol, we use skeletal muscle myosin II that we purify from chicken muscle from published methods (35). The myosin is stored in 25 mM KPO_4_, 0.6 M KCl, 10 mM EDTA, 1 mM DTT, pH 6.6, either directly after purification or from frozen stock. It is important to keep the myosin cold at all times to prevent loss of motor activity.
2. Add DTT to the thawed myosin solution to a final concentration of 10 mM. Use a 1 M DTT stock solution to limit the volume. The DTT reduces the thiol group on cysteines in the myosin protein, preparing them to react with the maleimide for labeling.
3. To prevent interference with the labeling reaction, remove the DTT by dialyzing overnight in 50 mM HEPES, 500 mM KCl, 1 mM EDTA, pH 7.6, at 4° C.
4. After dialysis, centrifuge the myosin solution at 100,000 x *g* at 4°C for 15 minutes to pellet any aggregates. Collect the supernatant.
5. Determine the concentration of myosin in the supernatant using a spectrophotometer. The myosin concentration may be estimated using an extinction coefficient at 280 nm of ∼148,000 M^−1^cm^−1^ for skeletal muscle myosin (38). This extinction coefficient assumes a “monomer” to consist of one heavy chain, one essential light chain, and one regulatory light chain.
6. Prepare fluorophore for labeling reaction by resuspending powdered fluorophore in dry DMSO to a concentration of 5 mM fluorophore. Note: Any fluorophore with a maleimide reactive group should work. We used Alexa 647 maleimide. Note: DMSO is hygroscopic and accumulates water. By “dry” DMSO, we refer to DMSO that has been stored in an environment to prevent water accumulation. To ensure the solvent is dry DMSO, we use a freshly opened ampule of DMSO for this step. Note: Do not put DMSO solutions on ice. DMSO has a melting point of 19°C (39), so it will need to be room temperature (or slightly warmer) to pipette. Caution: DMSO can penetrate gloves, so it is good practice to wear double nitrile gloves and take care to prevent spills.
7. To prepare the myosin for adding fluorophore, briefly take the myosin solution off ice and let warm to room temperature. Note: Do not leave myosin at room temperature for longer than necessary. Do this step after the fluorophore is suspended in DMSO.
8. As soon as the myosin solution is near room temperature, add the fluorophore solution to the myosin and mix rapidly, such that there is a 5:1 molar ratio of fluorophore: myosin. Place myosin solution on ice as soon as it is mixed with the fluorophore.
9. Incubate on ice for 1 hr, protected from light, then stop the reaction by adding DTT to a concentration of 1 mM. Use a 1 M DTT stock to minimize the volume added.
10. Remove unreacted fluorophores. We use either of two methods:

**Option 1:** Polymerize myosin by slowly diluting into F-buffer (10 mM Imidazole, 50 mM KCl, 1 mM MgCl_2_, 2 mM EGTA, 4mM ATP, pH 7.5). Incubate on ice 20 minutes. Spin at 8,000 x *g* in a refrigerated centrifuge at 4°C for 10 min to pellet. Discard supernatant containing free dye and resuspend/de-polymerize myosin with Myosin Storage Buffer (from Step 1). To remove any remaining dye, dialyze into excess Myosin Storage Buffer 3 times, for 15 minutes each using a 10 kD cutoff dialysis cup.
**Option 2:** Use a desalting column for buffer exchange into Myosin Storage Buffer (from Step 1) with the standard protocol from column manufacturer.
11. Estimate the fluorophore to protein ratio by measuring the concentration of myosin and fluorophore with a spectrophotomer, using extinction coefficients for myosin at 280 nm and for the fluorophore as reported by manufacturer. A ratio of 2-4 is typical.

The labeled myosin is now ready for snap freezing of aliquots in liquid nitrogen. Aliquots are stored at −80 °C for long term storage.

### Optional: Procedure to remove inactive myosin

Myosin can be used directly from a thawed aliquot in experiments, but some fraction of the myosin will be inactive. The following procedure describes how to remove inactive myosin. This is an optional step, but it can be crucial for reproducibility. Myosin aliquots used for approximately 3 days after thawing, stored at 4°C or on ice.

1. Polymerize unlabeled, phalloidin-stabilized actin:

a. Mix 10 µL 10x F-buffer, 1 µL 100 mM ATP, and ddH_2_0 such that the final volume (including actin in step b) will be 100 µL in a microcentrifuge tube.
b. Add actin monomer to a final concentration of 20 µM and pipette mix.
c. To stabilize actin filaments, add phalloidin to a final concentration of 6.7 µM and pipet mix. Caution: Phalloidin is a toxin (40). Avoid spills and contamination.
d. Incubate on ice for 20 minutes to allow actin to polymerize. After polymerization, the actin can be stored at 4°C for future spin-downs.
2. Pipet mix 10 µL phalloidin-stabilized actin, 3 µL 10x F-buffer, 0.3 µL 100 mM ATP, 6 µL 2M KCl, and ddH20 to a final volume of 30 µL (including volume associated with adding myosin in the next step) in a microcentrifuge tube on ice.
3. Add dimeric, labeled myosin from prepared stock to desired concentration maintaining a molar ratio of myosin to actin of ≤ 1:6.
4. Centrifuge the resulting solution cold (4° C) at 100,000 x *g* for 30 minutes. Inactive myosin will bind to actin filaments and remain bound. The inactive myosin and actin filaments will form a pellet during centrifugation.
5. Remove supernatant (containing active myosin) and store at 4° C for use in experiments.
6. Estimate the myosin concentration using the same spectrophotometric method to measure myosin concentration after labeling. Note: At the protein and ATP concentrations that remain in the supernatant, the 280 nm absorbance peak is typically obscured by the 260 nm ATP absorbance peak, so measuring the concentration through the absorbance associated with the fluorophore, or a relative fluorescence measurement is more accurate.

### Prepare oil-surfactant solution

Note: the oil-surfactant solution can also be purchased pre-mixed and ready to use.

1. Start with a clean, glass vial. The vial can be rinsed in water and ethanol to remove potential contaminants to the oil, which is a solvent for the surfactant. We use borosilicate glass vials, with a PTFE-lined cap.
2. Pick up a small amount of 008-fluorosurfactant using a pipet tip and deposit near the bottom of a glass vial. Measure the surfactant amount by weight using a balance with sub-milligram accuracy.
3. Pipet the appropriate volume of Novec-7500 oil to make a 2 wt% solution and close the vial. Note: The oil is volatile, so close the vial immediately after measuring.
4. Vortex at low speed to mix. The solution can now be stored at 4° C for a few weeks. Note: Oil-surfactant age and quality can influence both crowding and appearance of the actin. Actin that appears speckled, adhered to the surface, or does not crowd to the surface may indicate that a fresh oil surfactant solution is necessary. Storage under nitrogen gas can improve shelf life.

### Prepare sample chamber (Three options)

These are standard chambers that we use for preparing samples for microscopy.

### Option 1: Flowcell

A simple flowcell is a standard way of making microscope samples in many labs. For this protocol, it is particularly suited to the samples encased in emulsion drops. It can be difficult to achieve large, flat, fields of view with the surfactant surface coating described in this protocol, so for 2D samples options 2 and 3 are often more reproducible. The channel geometry also may cause uneven mixing of components added at later times, such as myosin.

1. Place 2 pieces of double-sided tape parallel to each other across the width of the microscope slide, to form the boundaries of the flowcell channel. The channel formed between the two pieces of tape should be approximately 2 mm in width. Note: Smaller channels help prevent emulsions from getting stuck in the beginning part of the channel. To make a good seal, use pieces of tape just longer than the width of the glass slide and trim after constructing the flowcell.
2. Place the coverslip over the tape channel. The coverslip should be perpendicular to the microscope slide. We prefer 30 mm x 40 mm coverslips for this purpose because provide a small overhang when placed on a slide (Fig. 4A).
3. To create a good seal and eliminate air pockets between the tape and the coverslip, press down on the area of the coverslip that is in contact with double-sided tape. Note: Be careful not to break the coverslip while pressing around the edges. Rubbing the area with a rounded object, such as the end of a capped marker, works well to press down the tape.
4. Using a razor blade, trim any excess tape off the microscope slide and coverglass, such that the only visible tape is within the sample channel. Note: Be careful completing this step as it is easy to accidentally break the coverslip while trying to remove the tape.

### Option 2: Cylinder on untreated coverslip

This option is suited for 2D planar samples. It is convenient for adding components at different times during microscopy experiments. This sample chamber should not be used for emulsions, which cream rather than sediment, rendering imaging difficult.

1. Rinse coverslip and glass cloning cylinder with water, ethanol, and water. Air dry.
2. Using a thin layer of 5-min epoxy, adhere the glass cloning cylinder to the coverslip (Fig. 2A). Note: It is helpful to apply force to the cylinder while the epoxy dries by placing a clean object on top of the cylinder. A small petri dish lid or the coverslip box can work well.

### Option 3: Cylinder on silane-coated coverslip

This option reproducibly results in a large, planar region in the center of the chamber and reduces bulk flow of the sample. This sample chamber should not be used for emulsions, which cream rather than sediment in buffer, rendering imaging difficult.

1. Place an octyl-silane treated coverslip into the bottom of a glass petri dish or any other open glass container that will fit into the UV-ozone cleaner.
2. Using forceps, place a 2 mm x 2 mm PTFE square (cut from a sheet) onto the coverslip.
3. Treat the coverslip with UV-ozone for 10 minutes. This process makes the glass hydrophilic in the exposed regions. The PTFE square may be reused in future experiments.
4. Carefully remove dish without disturbing the coverslip or Teflon square. Adhere a glass cloning cylinder to the coverslip using 5 minute epoxy (Fig. 2A). The cylinder should encircle the Teflon square, which may be removed with forceps after the epoxy dries.

### Assemble the actin network

Note: The following is for preparation of a 50 μL sample. For cylinder samples, a larger volume can reduce the effects of the meniscus and increase sample stability.

1. To a microcentrifuge tube, add:

a. 5 µL 10x F buffer
b. dd water necessary to bring final volume to 50 µL.
c. 1 µL 25 vol% betamercaptoethanol
d. 1 µL 225 mg/mL glucose
e. 1 µL mixture of glucose oxidase (135 mg/mL) and catalase (85,000 units/mL)
f. 1 μL 25 mM ATP
g. 10 µL 2 wt% 15 cp methylcellulose. Note: The methylcellulose stock solution is prepared by stirring at 4° C and can be stored at 4° C for several weeks. Methylcellulose can precipitate out of solution and should be re-prepared if there is visible precipitate or if actin crowding is poor. Pipet mix thoroughly.
2. Add cross-linking protein (∼0.1-1 cross-linker proteins for every 1 actin monomer) if desired. Pipet mix. Note: Cross-linking protein(s) may also be added after actin polymerization and allowed to bind for 10-20 minutes before adding myosin. In our experience, this is preferable for cross-linker concentrations that are high enough to bundle actin filaments—the bundles are more evenly and reproducibly distributed at the surfactant-water interface. The cylinder sample chamber is amenable to adding cross-linker after the actin polymerizes.
3. Add phalloidin, (∼1 phalloidin for every 10-3 actin monomers), if desired. Pipet mix.
4. In a separate tube, mix unlabeled and labeled actin to a total concentration of 1 µM in a ratio of 1 labeled actin monomer to 10 total actin monomers. Pipet mix. Note: The actin monomers will begin to polymerize upon addition to the sample solution. Adding the actin labeled and unlabeled actin separately can result in actin that appears speckled with dark and fluorescent regions along the contour of a single filament. We mix the actin monomer stocks to ensure uniformly labeled actin filaments. Actin purification and labeling is described in (31).
5. Optional: To make liquid crystals or other experiments using short actin filaments, add capping protein to the actin mix from step 4. The exact amount of capping protein depends on the active fraction of the protein and the length of actin filaments targeted, but typically around 1-10 mol% capping protein (with respect to the actin) is sufficient to create short filaments for liquid crystal experiments. Capping protein purification is described in (36).
6. To begin polymerizing actin, add the actin mix to the F-buffer solution and pipet mix.
7. The actin may be left to polymerize in the microcentrifuge tube at room temperature, and then added to the sample cell after 10-30 minutes. **Optional:** If preparing a cylinder sample, draw entire solution into a pipet tip so it is ready to pipet in step 4 of loading the cylinder sample chamber.
8. **Optional: Prepare emulsions**

a. Add 3.5 μL of oil-surfactant to a microcentrifuge tube. It is best to use a transparent, light colored or clear microcentrifuge tube for this step.
b. Without mixing, add 5 μL of the actin solution to the top of the oil-surfactant layer. Close the microcentrifuge tube.
c. While holding the top of the microcentrifuge tube, flick the bottom edge of the tube. This should initially create a foam with visible emulsions. Continue to flick the tube until the foam has no distinguishable features when you look at the tube backlit – this should create microemulsions.

Note: When preparing a sample in emulsions, a flowcell or other sample chamber should be ready before making the actin sample. The glass cylinder sample chamber does not work well for emulsions as they will cream to the air-surfactant solution interface rather than sediment to the coverglass surface.

### Load the sample chamber (Cylinder chambers)

1. Pipet 5 µL of oil-surfactant solution into the bottom of the glass cylinder chamber Note: Alternatively, samples can be made by passivating the surface with a supported lipid bilayer (11, 29, 30).
2. Tilt coverslip and slowly rotate such that the coverslip surface and lower cylinder are coated with the surfactant solution.
3. Remove excess oil-surfactant solution with pipet to get as thin of an oil layer as possible while not allowing the oil, which is volatile, to evaporate completely.
4. Immediately add the sample solution from step 7 (Assemble the actin network) to the chamber.
5. Cover the chamber with a small piece of Teflon tape to prevent evaporation and flow.

### Alternate: Load the sample chamber (Flowcell)

1. To prepare for loading the flowcell, mix a small amount of 5 min epoxy.
2. To place your sample in the flow cell, first pipette 1-3 μL of oil-surfactant solution into the sample channel of the flow cell, to create a small plug of solution that wets the channel.
3. Pipette the sample solution or emulsion suspension slowly into the entrance of the flowcell. The entrance is the same side that the oil-surfactant solution was added to. If the sample volume exceeds the flowcell volume, wick the sample through the flowcell by placing a kimwipe or filter paper at the other end of the channel. If the meniscus of the oil-surfactant solution has receded, resulting in an air gap, then first tilt the flowcell to remove the air gap at the entrance prior to adding the sample.
4. If necessary, add additional oil-surfactant solution until air is no longer visible in the channel or to push the sample (particularly for emulsions) further into the channel.
5. Seal each side of the channel with 5-minute epoxy (mixed right before loading the flowcell).

### Image and add myosin

1. Mount sample on microscope and start timelapse imaging of actin. If polymerizing in the sample chamber, allow polymerization for ∼30 minutes or until there is no longer visible filament lengthening or motion.
2. Optional: Add cross-linker.
3. Add myosin:

a. as a dimer pipetted to the top of the sample (no mixing is required because the myosin dimer will diffuse) or
b. as pre-polymerized filaments. Pipet mix approximately half of the total volume very slowly to avoid disturbing the crowded actin. Note: The ideal method for myosin addition to get the desired filament density or size at the coverslip surface may vary for different actin architectures and needs to be determined experimentally.
4. Start timelapse imaging.

## RESULTS

A broad range of actin assemblies can be formed in model systems using purified proteins by following the general strategy described in these methods, where actin is polymerized into filaments in the presence of accessory proteins that modify the assembly architecture (Fig. 1). When actin is polymerized into filaments without cross-linkers or capping protein, it forms entangled filament networks (Fig. 1A). Adding crosslinker to the entangled networks results in the formation of quasi 2D networks or networks of bundles (Fig 1B, C). Whether networks or networks of bundles form depends on the type and amount of cross-linker added to the actin network (e.g., α-actinin will form networks at low concentrations and networks of bundles at higher concentration as shown here). The cross-linker can also be added while the actin is polymerizing, although this can result in the formation of a 3D rather than quasi 2D network.

**Figure 1.**
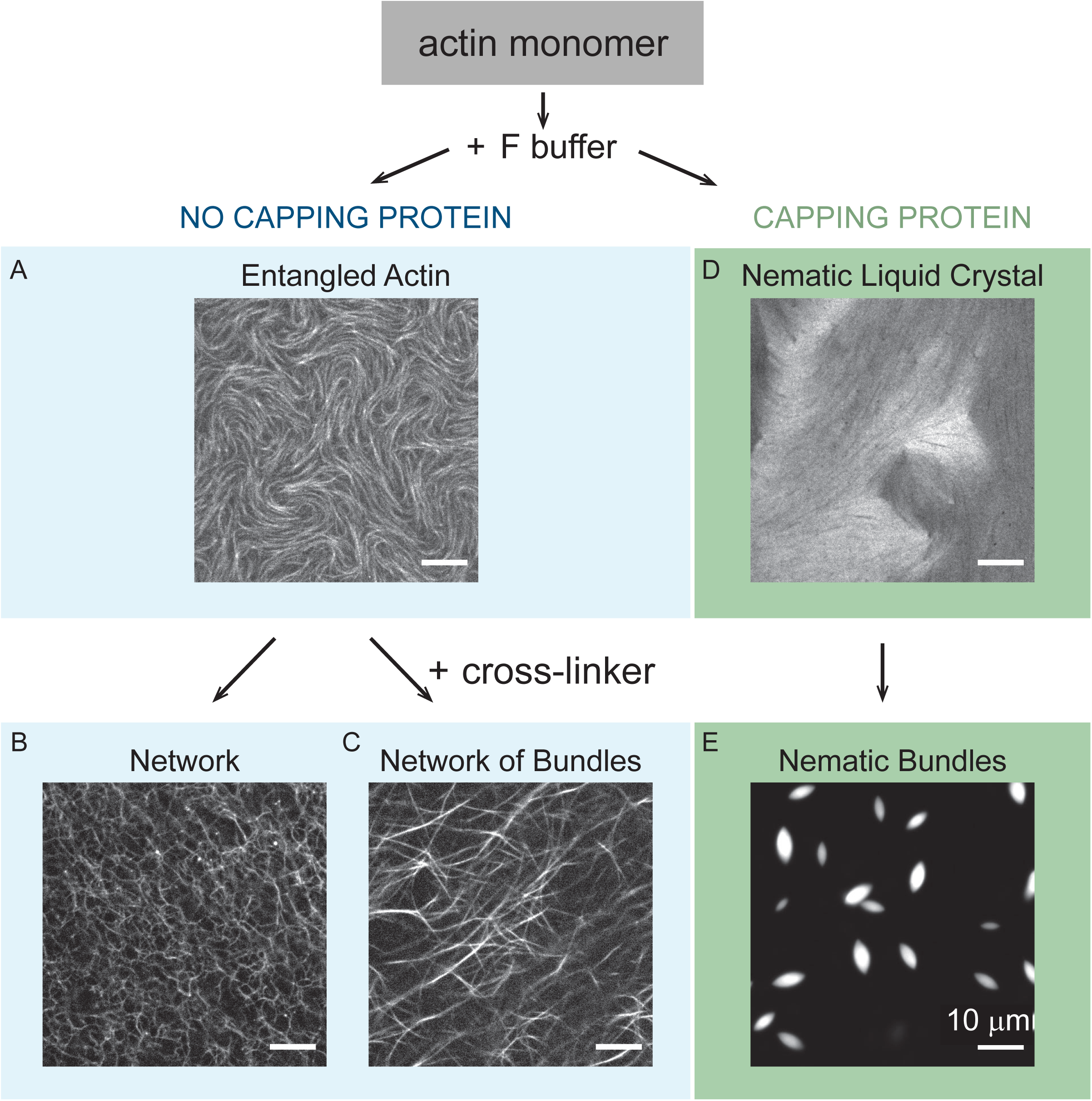
General method for forming model actin assemblies with various architectures. **(A)** Actin monomer is polymerized in F-buffer to form entangled actin filaments. **(B)** To form cross-linked networks, a small amount of network-forming cross-linker, such as 0.5 mol% alpha-actinin (pictured), is added after the entangled network is formed. Cross-linker can also be added during actin filament polymerization. **(C)** To form a network of bundles, an increased amount of network-forming cross-linker (1-10 mol%), or a bundle-forming cross-linker, such as 5 mol% fascin (pictured) is added to the entangled filament network. **(D)** To instead form an actin liquid crystal, actin monomer is initially polymerized in the presence of capping protein to form short actin filaments, which are crowded to the surface. **(E)** To form liquid crystal droplets of actin, 5 mol% of the cross-linker filamin is added. For all images, the scale bar is 10 μm.

To instead create liquid crystalline assemblies, actin is polymerized in the presence of capping protein, which limits actin filament growth (Fig 1. D, E). Filaments polymerized *in vitro* using the methods described in Fig. 1A result in filaments that are >10 µm long, which is similar to the persistence length of actin.). Filaments polymerized in the presence of capping protein can be below the resolution limit of the microscope (∼250 nm) in length, which is in a regime where actin can be thought of as a rigid rod. Short actin filaments, polymerized with capping protein and crowding agent present, form quasi 2D actin nematic liquid crystals (Fig. 1D). Insufficient capping protein can result in micron-sized filaments, which transition to elongated bundles instead forming an entangled network (11). By adding certain cross-linkers in the presence of nematic forming short actin filaments, nematic actin droplets, called tactoids, form instead of a 2D liquid crystal (Fig. 1 E). To our knowledge, cross-linked actin tactoids have been experimentally constructed only using the cross-linker filamin, although it is theoretically expected that actin tactoids could be formed with other cross-linkers.

Representative long, entangled actin networks formed in cylindrical sample chambers are shown in Fig. 2. The actin is uniformly distributed over the surface, with local alignment of filaments and differences in packing seen in intensity variations (Fig. 2B). Bright spots in the image indicate where filaments overlap, while small dark regions have little or no actin. The actin filaments will thermally fluctuate, leading to local intensity changes and visible bends in an image sequence or when viewing the data in real time acquisition. Adding myosin II (Fig. 2C, white spots) induces bending and reorganization of actin network (Fig. 2C). The time scale and extent of the myosin-driven network changes depends on the myosin concentration and sample conditions, such as ATP concentration. Representative results of myosin at a high enough local density to drive network contraction are shown in Fig. 2C and Movie 1 (image sequence of data in Fig. 2C). The extent of contraction can spatially vary over the sample if the local myosin concentration varies.

**Figure 2.**
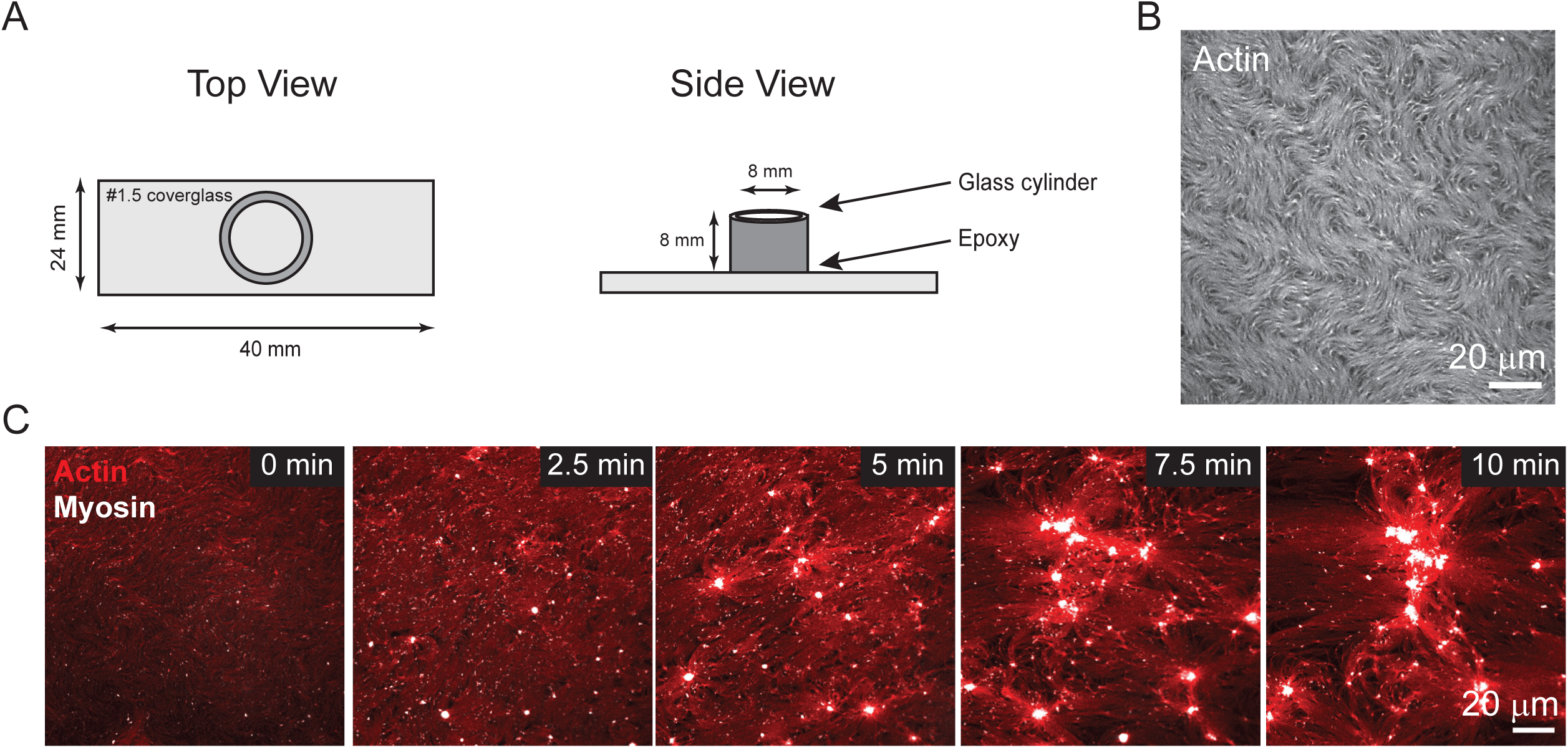
Representative results of an entangled actin network undergoing myosin-induced contraction. **(A)** Schematic of microscopy sample chamber. The sample is housed in a glass cylinder that is epoxied to a #1.5 coverglass. **(B)** Representative confocal fluorescence image of entangled actin network. Scale bar is 20 µm. **(C)** Representative results myosin-induced contraction of an entangled actin network (red). After mixing (t = 0 min), myosin puncta (white) bind to the network without inducing visible deformations. Myosin continue to accumulate in the network and begin to induce deformations (t = 2.5 min) and actin filament reorganization, until the myosin collects into aggregates at long times. Scale bar is 20 µm.

Representative results of cross-linked actin networks driven by myosin formed in cylindrical sample chambers are shown in Fig. 3. The details of initial network microstructure (such as the actin bundles seen in Fig. 3, 0 s) and the network remodeling is dependent on the amount and type of cross-linker. In cross-linked networks, the actin has coordinated motion over longer length scales (Fig. 3, Movie 2) than the more local contraction seen in entangled networks (Fig. 2C, Movie 1). The actin network motion reaches a steady state as the myosin clusters.

**Figure 3.**
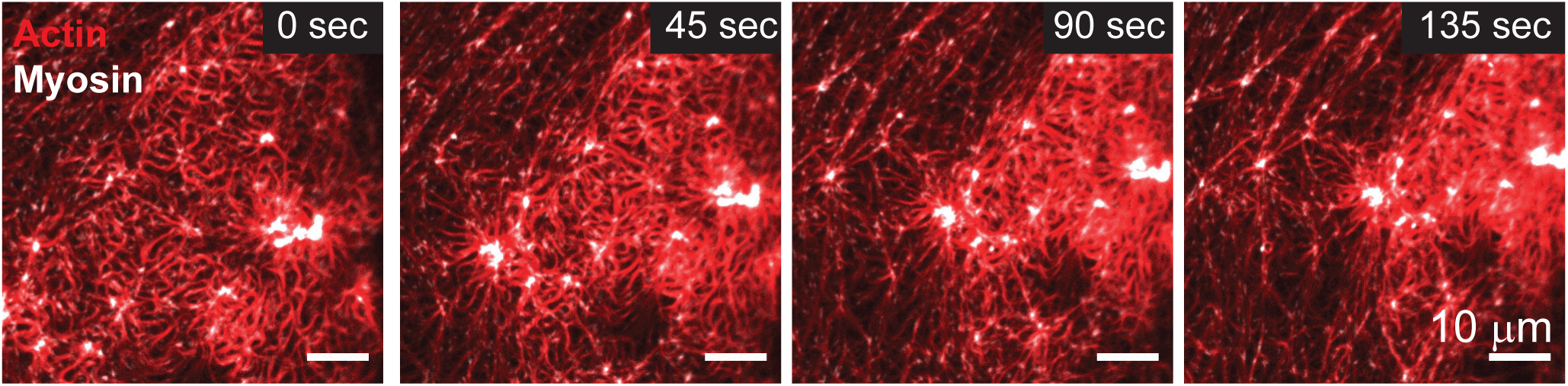
Representative results of a network of bundled actin filaments, cross-linked by 2.5 mol% undergoing myosin-induced contraction. After mixing (t = 0 sec), myosin puncta (white) bind to the cross-linked network without visible deformation. Myosin puncta continue to bind and coalesce, inducing visible contraction and reorganization of the cross-linked actin network (t = 45 sec). At long times, myosin-induced contraction ceases the deformation of the cross-linked network. Scale bar is 10 μm.

The sample chamber should be selected according to experimental needs. As an alternate to the cylinder sample chamber, we also describe a flowcell sample chamber (Fig. 4A). Representative data of an entangled actin network, formed in emulsions and imaged in a flowcell are shown in Fig.4B. Emulsions formed by this method vary in size. Large emulsions, with diameter larger than the gap between the coverglass and the slide glass (∼80 µm with the double stick tape spacer used here), are suitable for imaging actin networks (Fig. 4C & 4D). The actin can be polymerized in the emulsions (Fig. 4C) or in a microcentrifuge tube and later loaded into emulsion (Fig. 4D) yielding similar networks that are comparable to networks formed in cylindrical sample chambers (Fig. 2B).

**Figure 4.**
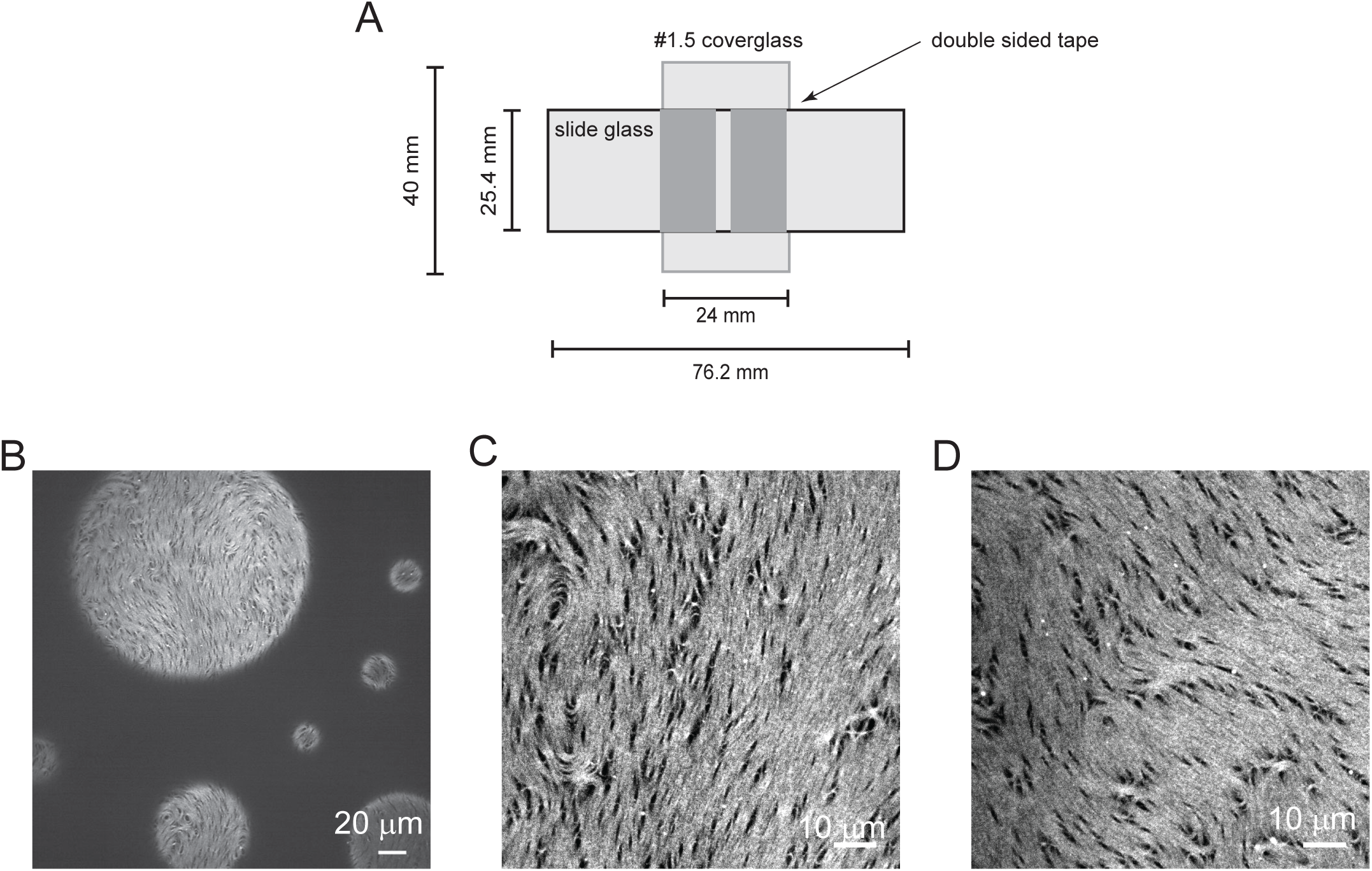
Alternative sample preparation methods. **(A)** Schematic of flowcell. The sample is wicked through a chamber between a microscope glass slide and a #1.5 coverglass with double-sided tape. **(B)** Representative confocal microscopy image of entangled actin networks in emulsions. Emulsions prepared by this method (flicking) have a range of sizes. Scale bar is 20 μm. **(C)** Representative image of entangled actin network polymerized in emulsion drops for approximately 15 min. Emulsion drops were added to the flowcell immediately after preparing. Scale bar is 10 μm. **(D)** Representative image of the entangled actin network polymerized in a microcentrifuge tube for 16 min before forming emulsion drops. The emulsion drops were added to the flowcell immediately after preparing. Scale bar is 10 μm.

## DISCUSSION

There are many considerations to producing reproducible actin assemblies. One of the most critical to producing reproducible, analyzable data is the coverglass surface of the sample chamber. The proteins in in vitro actin samples, particularly myosin, are extremely sticky and will adhere to untreated or poorly treated glass surfaces rendering a sample that is unusable (Fig 5A). We have discussed a standard way of prepping the surface to yield reproducible samples, through passivating a silanized coverslip with an oil-surfactant layer. However, this sample preparation has several variations depending on the resources available. The sample can be made on a silanized coverslip without UV/ozone treatment, or on a plain glass coverslip with an oil surfactant layer (sample cell option #2). These methods are simpler to prepare, but may result in the sample having an uneven oil surface (Fig. 5B), potentially making it challenging to find a large, flat region of the sample to analyze. The Teflon mask/uv/ozone method described can reduce this, resulting in one hemispherical surface, with a large planar region at the center. It is important to remove excess surfactant unintentionally end up as emulsions (Fig. 5C). A supported lipid bilayer (11, 29, 41) will generally produce larger, flatter surfaces than the surfactant solution described here. However, the supported lipid bilayer method has more preparation involved and is sensitive to contaminants and small environmental changes. When using bilayer, it is important to passivate the surface at the microscope, to avoid even small (∼1°C) changes in temperature. It is important to consider that supported bilayer can also be disturbed by certain components of the sample or protein storage buffers, even at extremely low concentrations, such as glycerol.

**Figure 5.**
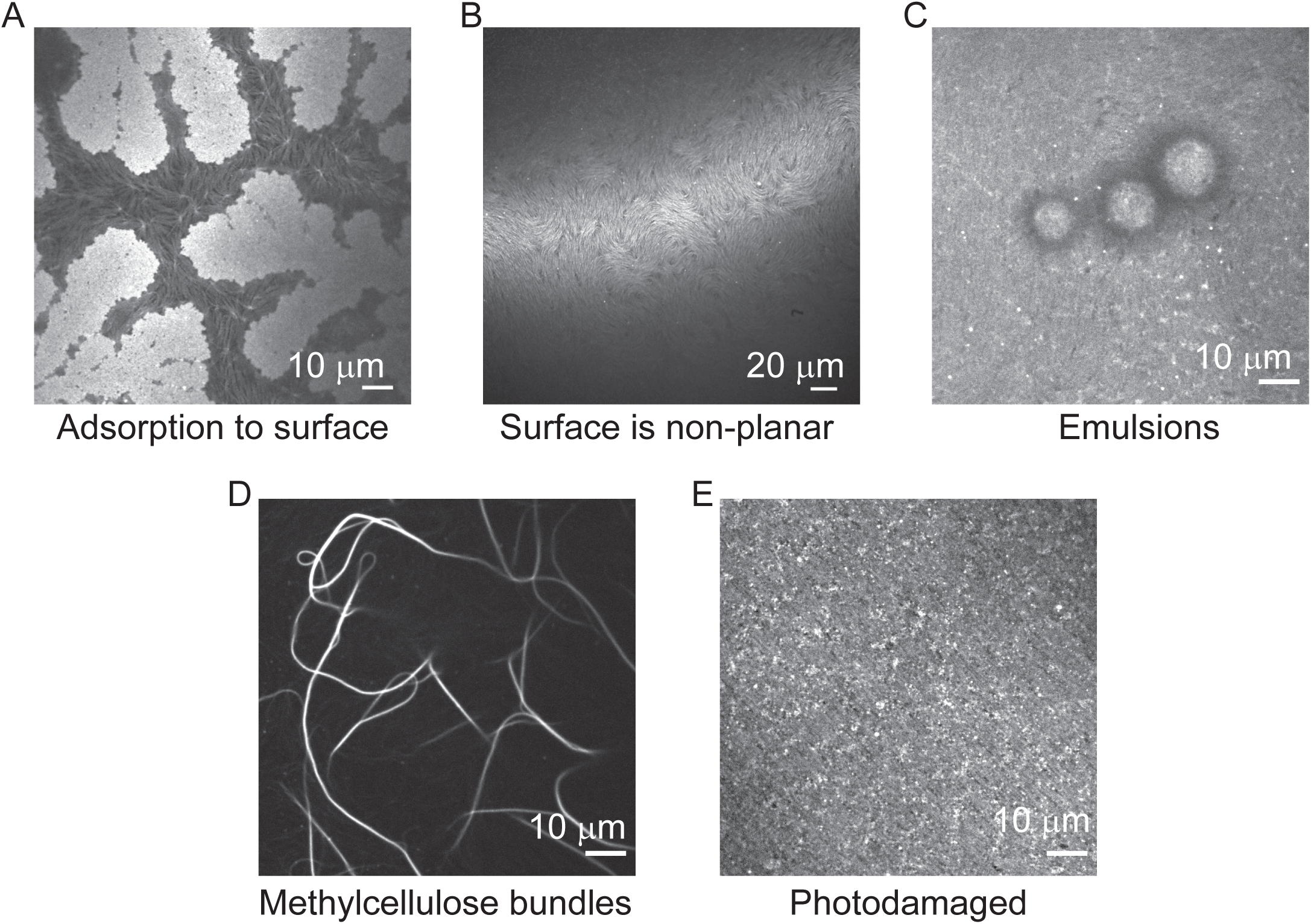
Common deviations from representative results. **(A)** Actin adsorbs to the surface, due to incomplete surface passivation. **(B)** The crowded actin layer is non-planar, caused by excess oil surfactant solution remaining after surface passivation. **(C)** Emulsions in the sample plane, indicating that the surfactant solution is not fully removed from the surface prior to adding the sample. **(D)** Actin filaments are bundled by methylcellulose. This occurs when methylcellulose concentration is greater than intended. **(E)** Actin is photodamaged. This occurs when light source intensity is too high, sample is imaged for extended period of time, or oxygen scavenging system components are not properly functioning.

The protein components of the model system are also key to reproducibility. The proteins that we regularly used have been purified either in house or commercially, then aliquoted and flash-frozen for long term storage at −80°C. We see no difference in house or commercially purified actin for the type of experiment described here. Once thawed, proteins are typically stable in their storage buffers on ice or at 4°C for approximately 3 days. The proteins can be photodamaged during microscopy imaging (Fig. 5E). Photodamaged actin can appear speckled and more connected than entangled actin. Photodamaged actin may adsorb to the surface and have reduced response to myosin motor activity. Lowering the intensity of the light source, the frequency of imaging, or using freshly prepared oxygen scavenging system reagents will reduce photodamage.

The depletion agent also is critical to achieving reproducible samples. Methylcellulose is used as a depletion agent in this protocol, for the purpose of crowding the actin into a quasi-2D layer at the coverglass. If the researcher would prefer to make a 3D network, then depletion agent is not necessary. Methylcellulose is a common polymer of choice in the actin community for crowding, but other depletion agents, such as polyethylene glycol which is commonly used to prepare microtubule samples, would be expected to produce similar results (8, 42, 43). Of note, methylcellulose is more soluble at lower temperatures and will precipitate out of solution at room temperature (44). To ensure the methylcellulose concentration is stable, we prepare a stock solution and keep it at 4°C. This solution is stable, when kept stirring (gently to avoid air bubbles) at 4°C. If left at 4°C not stirring (as in a refrigerator), it will need to be replaced with a fresh solution after a few months. When the methylcellulose concentration is higher than expected, either from an inhomogeneous solution or from error in pipetting (the solution is extremely viscous and can collect on the outside of a pipette tip), then methylcelloluse may bundle the actin filaments (Fig. 5D).

The details of how myosin is added to a sample depend on the type of actin assemblies being formed, the type of sample, and the equipment available. Typically, we try several methods and use the one that is the most reproducible for the sample needs. It is important to keep in mind that myosin II forms minifilaments in the sample buffer, which are polymers of several (up to hundreds) of myosin II dimers. When myosin is thawed from frozen stock, some fraction of the motor heads are inactive, and once polymerized into minifilaments, these inactive heads can bind to actin filaments but do not undergo the ATP driven powerstroke and unbind. Then, myosin filaments that contain inactive heads are still able to exhibit characteristics of motor activity, but may also cross-link actin filaments. It is important, then, to note whether the myosin has been “cleaned up” or if it is used as thawed frozen myosin.

If myosin is added as dimers, they quickly diffuse and reach the surface of the sample. If not mixed, there may be regions of the sample where more myosin activity is occurring. The sample must be gently pipette mixed when adding prepolymerized filaments to ensure the myosin uniformly incorporates in the sample. An advantage of prepolymerizing the filaments is control over the size of the myosin filaments, through protein concentration and salt concentration in the prepolymerization mixture (25). A disadvantage of prepolymerizing myosin filaments is that they can bind to actin in solution and take longer to reach the surface. Another consequence of binding to actin in solution is that a higher concentration of prepolymerized myosin might be needed to achieve the similar myosin surface concentrations as when myosin is added in dimer form.

## ACKNOWLEDGEMENTS

We thank Todd Thorenson, Mike Murrell, Jennifer Ross, and Patrick McCall, for useful discussions while developing the methods, as well as Gardel and Kovar labs (University of Chicago). M.A.C. and S.R. were partially supported by Clemson University’s College of Engineering Computing and Applied Sciences Undergraduate Research Opportunity Grants and V.J. A. was supported by the Clemson Biophysics REU under NSF Award #2349368 with funding from the DBI and EPSCoR programs. This work was also supported in part by the National Science Foundation EPSCoR Program under NSF Award #OIA-1655740, in part by and in part by the Clemson Creative Inquiry + Undergraduate Research program.

## REFERENCES

1. M. C. Marchetti, et al., Hydrodynamics of soft active matter. Rev. Mod. Phys. 85, 1143–1189 (2013).

2. D. Needleman, Z. Dogic, Active matter at the interface between materials science and cell biology. Nat Rev Mater 2, 17048 (2017).

3. M. Murrell, P. W. Oakes, M. Lenz, M. L. Gardel, Forcing cells into shape: the mechanics of actomyosin contractility. Nat Rev Mol Cell Biol 16, 486–498 (2015).

4. R. Cáceres, M. Abou-Ghali, J. Plastino, Reconstituting the actin cytoskeleton at or near surfaces in vitro. Biochimica et Biophysica Acta (BBA) - Molecular Cell Research 1853, 3006–3014 (2015).

5. F. Burla, Y. Mulla, B. E. Vos, A. Aufderhorst-Roberts, G. H. Koenderink, From mechanical resilience to active material properties in biopolymer networks. Nat Rev Phys 1, 249–263 (2019).

6. M. Bezanilla, A. S. Gladfelter, D. R. Kovar, W.-L. Lee, Cytoskeletal dynamics: A view from the membrane. Journal of Cell Biology 209, 329–337 (2015).

7. R. Alfaro-Aco, S. Petry, Building the Microtubule Cytoskeleton Piece by Piece. Journal of Biological Chemistry 290, 17154–17162 (2015).

8. T. Sanchez, D. T. N. Chen, S. J. DeCamp, M. Heymann, Z. Dogic, Spontaneous motion in hierarchically assembled active matter. Nature 491, 431–434 (2012).

9. P. J. Foster, S. Fürthauer, M. J. Shelley, D. J. Needleman, Active contraction of microtubule networks. eLife 4, e10837 (2015).

10. M. P. Murrell, M. L. Gardel, F-actin buckling coordinates contractility and severing in a biomimetic actomyosin cortex. Proc. Natl. Acad. Sci. U.S.A. 109, 20820–20825 (2012).

11. K. L. Weirich, et al., Liquid behavior of cross-linked actin bundles. Proc. Natl. Acad. Sci. U.S.A. 114, 2131–2136 (2017).

12. J. Alvarado, M. Sheinman, A. Sharma, F. C. MacKintosh, G. H. Koenderink, Molecular motors robustly drive active gels to a critically connected state. Nature Phys 9, 591–597 (2013).

13. F. C. Keber, et al., Topology and dynamics of active nematic vesicles. Science 345, 1135–1139 (2014).

14. R. Zhang, N. Kumar, J. L. Ross, M. L. Gardel, J. J. De Pablo, Interplay of structure, elasticity, and dynamics in actin-based nematic materials. Proc. Natl. Acad. Sci. U.S.A. 115 (2018).

15. V. Schaller, C. Weber, C. Semmrich, E. Frey, A. R. Bausch, Polar patterns of driven filaments. Nature 467, 73–77 (2010).

16. L. Blanchoin, R. Boujemaa-Paterski, C. Sykes, J. Plastino, Actin Dynamics, Architecture, and Mechanics in Cell Motility. Physiological Reviews 94, 235–263 (2014).

17. P. M. Bendix, et al., A Quantitative Analysis of Contractility in Active Cytoskeletal Protein Networks. Biophysical Journal 94, 3126–3136 (2008).

18. S. Köhler, V. Schaller, A. R. Bausch, Structure formation in active networks. Nature Mater 10, 462–468 (2011).

19. F. Backouche, L. Haviv, D. Groswasser, A. Bernheim-Groswasser, Active gels: dynamics of patterning and self-organization. Phys. Biol. 3, 264–273 (2006).

20. T. M. Svitkina, Actin Cell Cortex: Structure and Molecular Organization. Trends in Cell Biology 30, 556–565 (2020).

21. A.-C. Reymann, et al., Actin Network Architecture Can Determine Myosin Motor Activity. Science 336, 1310–1314 (2012).

22. A.-C. Reymann, et al., Nucleation geometry governs ordered actin networks structures. Nature Mater 9, 827–832 (2010).

23. S. K. Vogel, Z. Petrasek, F. Heinemann, P. Schwille, Myosin motors fragment and compact membrane-bound actin filaments. eLife 2, e00116 (2013).

24. N. L. Liebe, et al., Bioinspired Membrane Interfaces: Controlling Actomyosin Architecture and Contractility. ACS Appl. Mater. Interfaces 15, 11586–11598 (2023).

25. T. Thoresen, M. Lenz, M. L. Gardel, Thick Filament Length and Isoform Composition Determine Self-Organized Contractile Units in Actomyosin Bundles. Biophysical Journal 104, 655–665 (2013).

26. M. R. Stachowiak, et al., Self-Organization of Myosin II in Reconstituted Actomyosin Bundles. Biophysical Journal 103, 1265–1274 (2012).

27. J. D. Winkelman, C. G. Bilancia, M. Peifer, D. R. Kovar, Ena/VASP Enabled is a highly processive actin polymerase tailored to self-assemble parallel-bundled F-actin networks with Fascin. Proc. Natl. Acad. Sci. U.S.A. 111, 4121–4126 (2014).

28. S. Stam, et al., Filament rigidity and connectivity tune the deformation modes of active biopolymer networks. Proc. Natl. Acad. Sci. U.S.A. 114 (2017).

29. K. L. Weirich, S. Stam, E. Munro, M. L. Gardel, Actin bundle architecture and mechanics regulate myosin II force generation. Biophysical Journal 120, 1957–1970 (2021).

30. M. Scholz, K. L. Weirich, M. L. Gardel, A. R. Dinner, Tuning molecular motor transport through cytoskeletal filament network organization. Soft Matter 16, 2135–2140 (2020).

31. J. A. Spudich, S. Watt, The Regulation of Rabbit Skeletal Muscle Contraction. Journal of Biological Chemistry 246, 4866–4871 (1971).

32. Y. Li, et al., The F-actin bundler α-actinin Ain1 is tailored for ring assembly and constriction during cytokinesis in fission yeast. MBoC 27, 1821–1833 (2016).

33. C. T. Skau, D. R. Kovar, Fimbrin and Tropomyosin Competition Regulates Endocytosis and Cytokinesis Kinetics in Fission Yeast. Current Biology 20, 1415–1422 (2010).

34. D. Vignjevic, et al., Role of fascin in filopodial protrusion. The Journal of Cell Biology 174, 863–875 (2006).

35. S. S. Margossian, S. Lowey, “[7] Preparation of myosin and its subfragments from rabbit skeletal muscle” in Methods in Enzymology, (Elsevier, 1982), pp. 55–71.

36. S. Palmgren, P. J. Ojala, M. A. Wear, J. A. Cooper, P. Lappalainen, Interactions with PIP2, ADP-actin monomers, and capping protein regulate the activity and localization of yeast twinfilin. The Journal of Cell Biology 155, 251–260 (2001).

37. Craig, S. W., C. L. Lancashire, and J. A. Cooper. 1982. Preparation of smooth muscle alpha-actinin. Methods Enzymol. 85:316–321.

38. A. B. Verkhovsky, G. G. Borisy, Non-sarcomeric mode of myosin II organization in the fibroblast lamellum. The Journal of cell biology 123, 637–652 (1993).

39. R. G. LeBel, D. A. I. Goring, Density, Viscosity, Refractive Index, and Hygroscopicity of Mixtures of Water and Dimethyl Sulfoxide. J. Chem. Eng. Data 7, 100–101 (1962).

40. A. M. Lengsfeld, I. LOWt, T. WIELANDt, P. Dancker, W. Hasselbach, Interaction of Phalloidin with Actin. Proc. Nat. Acad. Sci. USA (1974).

41. K. L. Weirich, J. N. Israelachvili, D. K. Fygenson, Bilayer Edges Catalyze Supported Lipid Bilayer Formation. Biophysical Journal 98, 85–92 (2010).

42. A. J. Tan, et al., Topological chaos in active nematics. Nat. Phys. 15, 1033–1039 (2019).

43. V. Nasirimarekani, T. Strübing, A. Vilfan, I. Guido, Tuning the Properties of Active Microtubule Networks by Depletion Forces. Langmuir 37, 7919–7927 (2021).

44. E. Heymann, Studies on sol-gel transformations. I. The inverse sol-gel transformation of methylcellulose in water. Trans. Faraday Soc. 31, 846 (1935).

